# Natural history of *Camponotus renggeri* and *Camponotus rufipes* (Hymenoptera: Formicidae) in an Atlantic Forest reserve, Brazil

**DOI:** 10.1101/2021.06.21.449232

**Authors:** Miguel Piovesana Pereira-Romeiro, Gabriel Tofanelo Vanin, Marianne Azevedo-Silva, Gustavo Maruyama Mori

## Abstract

Widespread species face a wide variety of environmental challenges and their morphology, behavior, and natural history may change across their range. However, not rarely, natural history research is restricted to one or few locations. That is the case for *Camponotus renggeri* and *C. rufipes*. Both species occur across South America in different ecosystems, but most research on these species is restricted to the Brazilian savanna, known as Cerrado. Here, we describe the foraging area, nesting habits, and activity schedule of *C. renggeri* and *C. rufipes* in an Atlantic Forest reserve in SE Brazil. *C. renggeri* foraged exclusively during nighttime and *C. rufipes* remained active throughout the day, but with little intensity during daylight hours. Most nests of both species were composed of dry straw, and average foraging areas were 0.91 m^2^ for *C. renggeri* and 1.79 m^2^ for *C. rufipes*. Inferred intraspecific foraging areas of nearby nests overlapped, especially for *C. renggeri*. Our findings reinforces the importance of natural history and what it adds to our knowledge on the ecology and behavior of *C. renggeri* and *C. rufipes* in Atlantic Forest.

---

The closely related carpenter ants *Camponotus renggeri* (Emery 1894) and *C. rufipes* (Fabricius 1775) have been extensively studied in terms of physiology (Takahashi-Del-Bianco *et al*. 1998; Galizia *et al*. 1999), natural history (Ronque *et al*. 2016; Ronque *et al*. 2018) and genetics (Azevedo-Silva *et al*. 2015; Ronque *et al*. 2016; De Aguiar *et al*. 2017; Azevedo-Silva *et al*. 2020). These species are morphologically similar, being distinguished mainly by the brightness of their tegument and by the color of their legs. Both species are commonly found on vegetation, where they feed on liquid resources from extrafloral nectaries and trophobionts (Oliveira & Freitas 2004; Ronque *et al*. 2018). Despite similarities in distribution and interactions with plants, these species differ with respect to nest architecture, foraging areas, and breeding systems. *C. renggeri* occupies nests built in dead tree trunks or underground, whereas *C. rufipes* may also construct nests out of dry straw (Ronque *et al*. 2016; Ronque *et al*. 2018; Azevedo-Silva *et al*. 2020). Both species occur across South America in different ecosystems (Janicki *et al*. 2016), but most research on these species is restricted to the Brazilian savanna, known as Cerrado. This ecosystem is characterized by an annual rainfall of 800 - 2,000 mm, with a very strong dry season during winter and fire-adapted flora (Oliveira-Filho & Ratter 2002).

*C. rufipes* and *C. renggeri* also occur in the Atlantic Forest, which, like Cerrado, is a conservation hotspot (Myers 2000), and presents highly diverse vegetation physiognomies. However, these biomes are markedly differentiated by seasonal precipitation and plant communities. Compared to Cerrado, Atlantic Forest presents wider latitudinal, longitudinal and altitudinal ranges and it is usually found in areas with higher precipitation, with an annual rainfall of 1,100 - 3,600 mm, specially in coastal regions (Oliveira-Filho & Fontes 2000). In this environment, natural history traits like foraging area, nesting habits, and activity schedule of these species remain largely undescribed. Given animal behavior is shaped by genetic, morphology, physiology, which are, in turn, influenced by the environment (Goodenough et al. 2009), it is expected that the natural history of these species may also vary in these distinct biomes.

The main goal of this study was to investigate the natural history of *C. renggeri* and *C. rufipes* in a distinct environment, the Atlantic Forest. Specifically, we aimed to describe their foraging area, nesting habits, and activity schedule. Our study is the first step towards a broader understanding of how the ecology and behavior of tropical ant species may vary across different ecosystems.

Fieldwork was carried out in an Atlantic Forest fragment surrounded by the sea and urban environment, at Xixová-Japuí State Park, São Vicente, São Paulo in Southeast Brazil (23°59’33.3” S, 46°23’21.7” W). The Park is dominated by secondary ombrophilous forests, and some areas are at initial successional stage, with abundant herbaceous plants and low canopy cover (Figure 1a). The region has a typically hot and humid climate, with mean annual temperature of 23.6°C and mean annual precipitation of 2,500 mm, being classified as tropical rainforest climate (Peel et al. 2007), which is environmentally contrasting to Cerrado areas.

**Figure 1.**
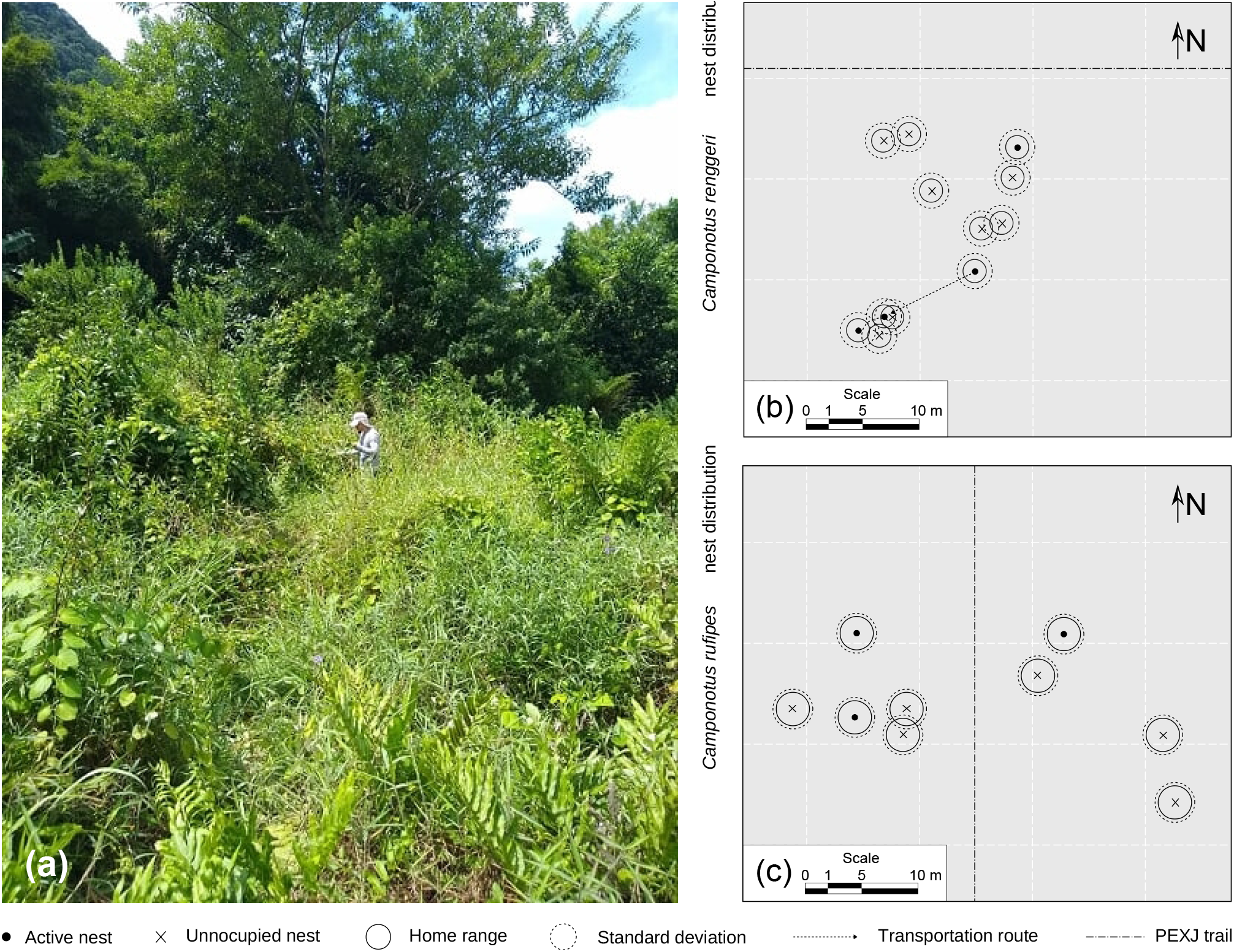
(a) General view of the Atlantic Forest reserve physiognomy in which the study was carried out. Distribution map of (b) *Camponotus renggeri* and (c) *C. rufipes* nests within plots where these species were found. Each spot represents a tagged nest, darker circles around them represent the mean foraging range for each species in the Atlantic Forest; outlined circles represent the standard deviation. Dark dots represent nests which were occupied by the end of the research, and X’s represent unoccupied nests by the end of the research. The larger dotted lines indicate trails of the at Xixová-Japuí State Park (PEXJ), in Southeast Brazil. The smaller dotted line represents an observed transportation route between nests.

Observations were performed from February to April 2019 and from May to July 2019. To investigate the co-occurrence of species, the study area was divided into ten plots (400 m^2^ each, 4000 m^2^ in total), where ants were searched for twenty minutes in each plot. In plots where they were found, we used sardine baits to find *C. renggeri* and *C. rufipes* nests. Additionally, we opportunistically found nests along the trails which were used for further analyses. Out of all the nests found, three to five of each species were tagged and characterized for activity schedule, nest architecture and foraging area.

Daily activity schedule was described by the total number of workers entering and exiting nests during sessions of 30 min, every 2 h, over a period of 24 h. To estimate foraging area, four *C. renggeri* and three *C. rufipes* nests were observed for three 5 hours 20 minutes observation periods in three different days, during their peak activity period (see results), summing a maximum of 16 h of observation for each species. Workers exiting the nest were followed until they returned, and the furthest distance was recorded. These points were tagged and had their distance to the nest measured and relative geographical location determined with a compass. Foraging area was estimated using minimum convex polygons generated with the package *adehabitatHR* (Calenge 2006) in R (R Core Team 2020). Due to low nest persistence, all observations of *C. renggeri* were carried out in the dry season, whereas observations of *C. rufipes* were performed during rainy season. During fieldwork, additional natural history records were made for both species using video, photographs, and notes (total of 32 h of observations for each species). We obtained hourly averages of air temperature and humidity data for each day from Brazil’s National Institute for Space Research (INPE) automatic weather stations database.

Nests were found in only three plots, and none of them had nests of both species. A total of 15 *C. renggeri* (12 nests in the same plot, one in a second plot and two nests opportunistically found outside of the plots) and nine *C. rufipes* nests (all within the same plot) were identified and tagged. Within a six-month interval (February to July 2019), out of the total, eight (53,3%) *C. renggeri* and six (50%) *C. rufipes* nests were unoccupied, indicating low nest persistence (Figure 1b, c). All *C. renggeri* nests found were built using dry straw (N = 15 nests), and some could have additional materials in their structure, such as freshly fallen leaves (N = 3) or dry leaves (N = 10). For *C. rufipes*, eight nests were built using dry straw as the main structural component, with four of them having dry leaves as additional material. A single underground nest belonging to this species was found. Many of these nests were suspended above ground, sustained by twigs (see Figure S1).

For the observed activity schedule, the cumulative number of nests entrances/exits per 30-minutes interval for both species indicated that the period of peak activity was nocturnal, increasing after sunset (18:00 h) and decreasing before sunrise (06:00 h) (Figure 2). Three *C. rufipes* nests were observed during hot/rainy season, with an average of 27.67 (SD: ± 8.38) workers sampled per nest. The observed foraging area for this species was 1.79 m^2^ (SD: ± 0.79 m^2^; min.: 1 m^2^; max.: 2.87 m^2^). For *C. renggeri*, four nests were observed in dry/cold season, with an average of 12.5 (SD: ± 3.84) workers sampled, resulting in an average foraging area of 0.91 m^2^ (SD: ± 1.10 m^2^; min.: 0.13 m^2^; max.: 2.81 m^2^). Although we were unable to find nests of both species in the same plot, and foraging area was small for the observed nests, we recorded major workers of *C. renggeri* and *C. rufipes* engaging in aggressive behavior towards each other when crossing paths, resulting in the decapitation of *C. rufipes* (Video S1). These observations suggested that these species may share foraging areas and competition may occur between them.

**Figure 2.**
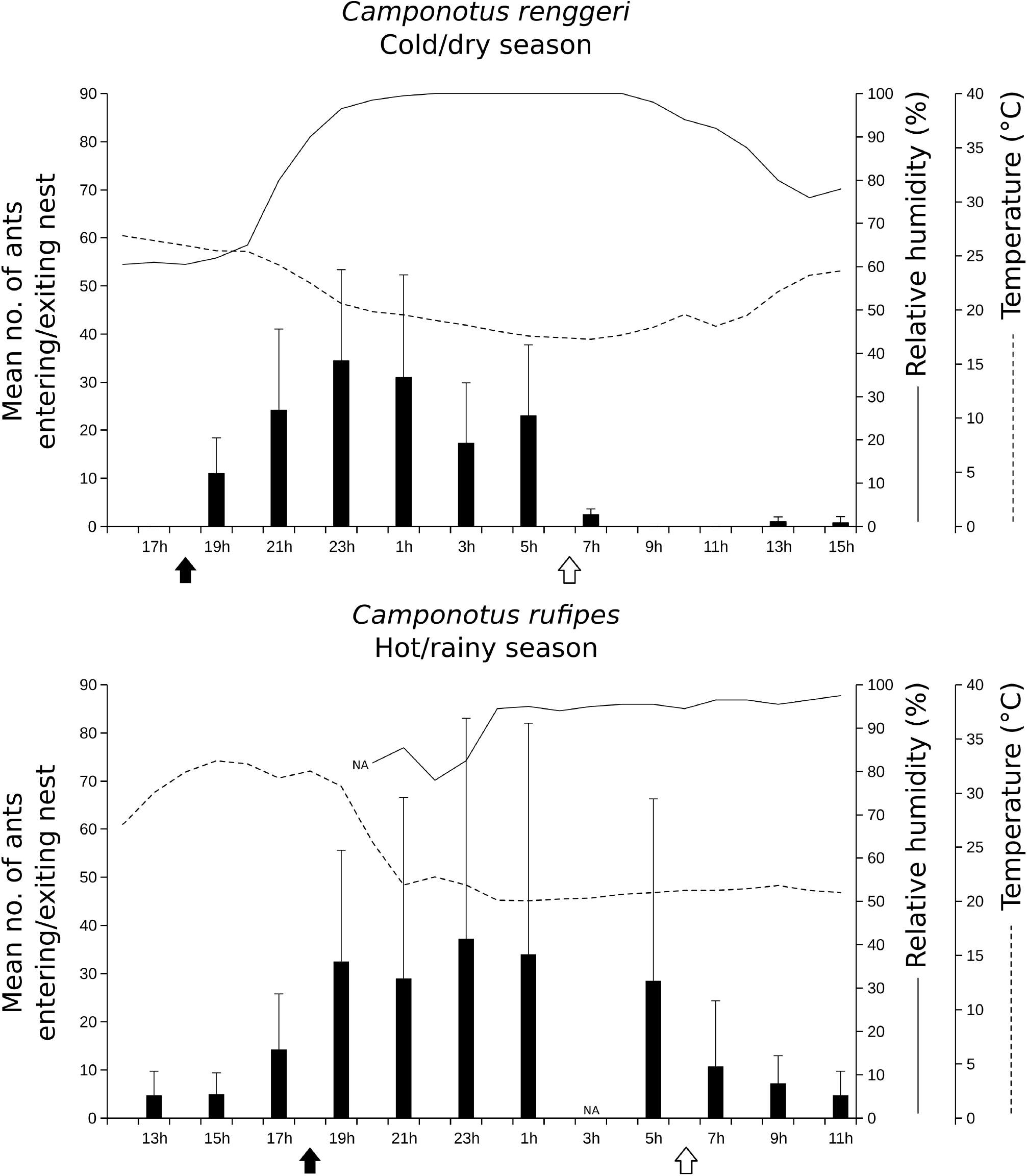
Daily activity of *Camponotus renggeri* (N = 4) and *C. rufipes* (N = 3) colonies in an Atlantic Forest reserve, southeastern Brazil. Daily activity was calculated by the cumulative number of ants leaving/entering the nests per 30-minutes interval (Mean ± SD). Air temperature and humidity for each day were collected from Brazil’s National Institute for Space Research (INPE) database (Mean), and humidity data was missing from some hours of the day. The arrows indicate sunrise (white) and sunset (black). NA stands for not available.

Our findings in the Atlantic Forest differ from those reported for other ecosystems, indicating that environment may play an important role in shaping *C. renggeri* and *C. rufipes* behavior. All but one *C. rufipes* nests characterized in this study were made of dry straw, similarly to the observed in the Cerrado and in the Argentinian “Chaco” (Weidenmüller *et al*. 2009; Ronque *et al*. 2018). Conversely, *C. renggeri* also presented this same nest architecture, differing from the nests in dead trunks reported for this species in Cerrado (Ronque *et al*. 2018), whereas no nests of such type were found in the Atlantic Forest. It is known that vegetation may play a role in regulating availability of resources for nesting in ants (Ronque *et al*. 2018). The Atlantic Forest fragment where this research was conducted is dominated by herbaceous plants, thus, dead trunks are not as abundant as leaf litter. This may explain the prevalence of nests made of dry straw over other nest architectures (Silvestre *et al*. 2003). The difference found for *C. renggeri* nests between different ecosystems indicates that this species is able to modify its nesting behavior in response to resource availability in different environments. Additionally, our findings expand the repertoire of nesting strategies reported for the *Camponotus* genus (Pfeiffer & Linsenmair 2000; Santos *et al*. 2005; Tschinkel 2005; Yamamoto & Del-Claro 2008; Santos & Del-Claro 2009).

*Camponotus rufipes* and *C. renggeri* presented smaller foraging area compared to other *Camponotus* species (138.38m^2^ for *C. sericeiventris* (Yamamoto & Del-Claro 2008), ∼230m^2^ for *C. cruentatus* (Alsina *et al*. 1988), ∼1700m^2^ for *C. leydigi* (Soares & Oliveira 2021), ∼8000m^2^ for *C. gigas* (Pfeiffer & Linsenmair 2000)). Additionally, despite the impracticability of a direct comparison due to slight methodological differences, we observed smaller foraging areas than the smallest ones reported for these species in Cerrado (*C. renggeri*: 2.78 ± 1.76 m^2^; *C. rufipes*: 4.55 ± 3.41 m^2^, Ronque *et al*. 2018). Also, in the Cerrado *sensu stricto, C. rufipes* average foraging area nearly doubled during rainy season (hot/rainy season: 9.83 ± 2.57 m^2^, dry/cold: 4.55 ± 3.41 m^2^), whereas *C. renggeri* foraging area was not affected by different seasons (hot/rainy season: 2.98 ± 1.28 m^2^, dry/cold: 2.78 ± 1.76 m^2^) (Ronque *et al*. 2018). Given our observations occurred in different seasons for *C. rufipes* and *C. renggeri*, it is unclear whether the foraging area estimated for *C. rufipes* is indeed larger than *C. renggeri* in the Atlantic Forest or if it is an effect of seasonality. However, the differences reported between biomes may be related to many factors, such as temperature, humidity, colony size, competition, and food availability (Breed *et al*. 1990; Gordon 1995; McGlynn *et al*. 2003; Ronque *et al*. 2018).

Beyond seasonal changes in foraging, daily changes in temperature, humidity and light may also influence a colony’s activity (Hölldobler & Wilson 1990). We showed that *C. renggeri* foraged exclusively during night in the cold/dry season, whereas *C. rufipes* remained active throughout day and night, but had predominant activity during nighttime in the hot/rainy season. Similar patterns were reported for this species in Cerrado, with *C. renggeri* activity being negatively affected by temperature and positively by humidity, whereas *C. rufipes* was not affected by these environmental conditions (Ronque *et al*. 2018). Indeed, nocturnal behavior is common among tropical ant species, presumably because it allows them to avoid high temperatures and low humidity during the day (Pfeiffer & Linsenmair 2000; Ronque *et al*. 2018). Additionally, temporal segregation can be a result of competition and resource sharing, particularly in exudate-feeding *Camponotus* species (Del-Claro & Oliveira 1999; Oliveira *et al*. 1999; Ronque *et al*. 2018). Accordingly, we recorded an aggressive encounter of major workers of *C. renggeri* and *C. rufipes*, which further suggest an antagonistic relationship between these species. However, because these ants can combine exudate with animal prey in their diets (Oliveira & Freitas 2004; Ronque *et al*. 2018), other factors besides competition and resource sharing could explain the observed difference in daily activity and the observed spatial partition between *C. renggeri* and *C. rufipes*.

This work focused on the description of natural history traits of *C. renggeri* and *C. rufipes* in the Atlantic Forest, an ecosystem where the ecology of these species is poorly explored. Our findings suggest that the behavior of studied ants may vary in different environments, shedding new light on the natural history of *Camponotus* species in tropical ecosystems. Identifying and understanding such variation is crucial to elaborate more complex and robust questions regarding ant ecology. Finally, we expect that our work encourages further investigation on ant natural history, which would be important to increase our knowledge on tropical systems.

## ACKNOWLEDGMENTS

We thank P. S. Oliveira, M. U. V. Ronque, and two anonymous reviewers for their valuable comments and suggestions that improved this paper. Also, the authors thank Instituto Florestal (COTEC 260108–010.270/2018), Instituto Chico Mendes de Conservação da Biodiversidade (SISBIO 66314-1), and Conselho de Gestão do Patrimônio Genético (A025CBF) for sampling and genetic resources access licenses. This research was funded by the Conselho Nacional de Desenvolvimento Científico e Tecnológico (CNPq – PIBIC/Unesp #48498, #48010, and #54099) and Fundação de Amparo à Pesquisa do Estado de São Paulo (FAPESP 2017/18291-2; 2019/12646-9; 2020/15636-1). This study was financed in part by Coordenação de Aperfeiçoamento de Pessoal de Nível Superior – CAPES (Finance Code 001).

## SUPPORTING INFORMATION

Additional Supporting Information may be found online in the Supporting Information section at the end of the article.

**Figure S1.**
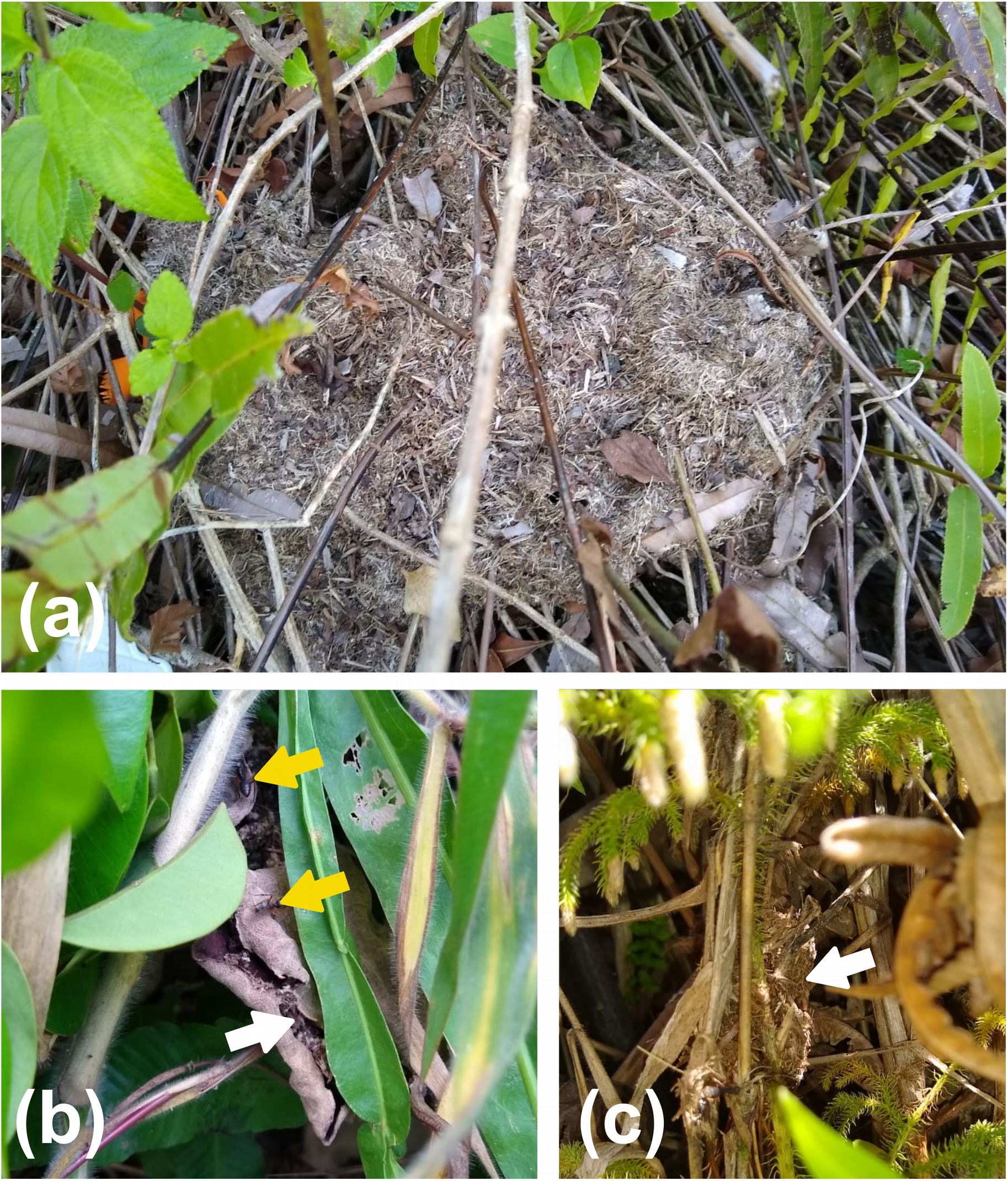
*Camponotus renggeri* and *C. rufipes* nests found in an Atlantic Forest Reserve, southeastern Brazil. Note the presence of whole leaves and intertwining branches in the architecture. The figure shows (a) a large nest of *C. renggeri*, (b) a small nest of *C. renggeri* inside the cavity formed by a twisted leaf with workers around it, and (c) small nest of *C. rufipes* entangled in vegetation. Yellow arrows indicate workers and white arrows indicate the nests.

**Video S1**. Workers of *Camponotus renggeri* and *C. rufipes* engaging in aggressive behavior against each other in an Atlantic Forest reserve.

